# Tissue Kallikrein 1 cleaves complement factor C3 and activates the alternative complement pathway

**DOI:** 10.1101/2025.09.08.674260

**Authors:** SE Sartain, PM Jacobi, V Qian, B Loredo, M Ranjan, J Bueno, S Bhar, A Li, A Yee

## Abstract

Hematopoietic stem cell transplant-associated thrombotic microangiopathy (HSCT-TMA), characterized by microvascular endothelial damage and severe renal injury, negatively affects HSCT survivorship with high mortality and long-term renal morbidity. After HSCT, innate immunity and inflammation are often dysregulated. The alternative complement pathway (AP) of the innate immune system is overactivated in HSCT-TMA, but the mechanisms of its initiation are poorly described. Complement component C3 of the AP can be cleaved by proteases outside of the AP. Because mRNA expression of tissue kallikrein 1 (KLK1), an inflammatory serine protease that produces kinins, has been found to be markedly elevated in the renal endothelium of inflamed mice, we hypothesized that increased KLK1 activity during inflammation contributes to AP overactivation and endothelial injury in HSCT-TMA. We assessed AP activation, KLK1 activity, endothelial injury, and renal function in HSCT-TMA experimental models and disease settings and investigated C3 cleavage and AP activation by KLK1. We found that patients with HSCT-TMA had significantly increased AP activation and decreased KLK1 inactivation at TMA diagnosis compared to pre-HSCT. Mice challenged with HSCT-TMA triggers cyclosporine A (CsA) and lipopolysaccharide (LPS) exhibited increased AP activation, renal endothelial injury, and impaired renal function in the setting of decreased KLK1 inactivation. We further demonstrated that KLK1 cleaved AP component C3 to C3b that functionally activated the AP. Our data indicate a noncanonical mechanism for AP activation by KLK1 in settings of HSCT-TMA.

**Key Points:**

- Tissue kallikrein (KLK1) functionally cleaves complement factor C3.
- Reduced KLK1 inhibition is associated with activation of the alternative pathway of complement.

Graphical Abstract.
HSCT-TMA triggers (e.g. inflammation, immunosuppression) upregulate expression of the inflammatory serine protease KLK1, which cleaves C3 to initiate AP activation and amplification, leading to renal endothelial injury. Created with BioRender.com under its Academic License Terms with Baylor College of Medicine.

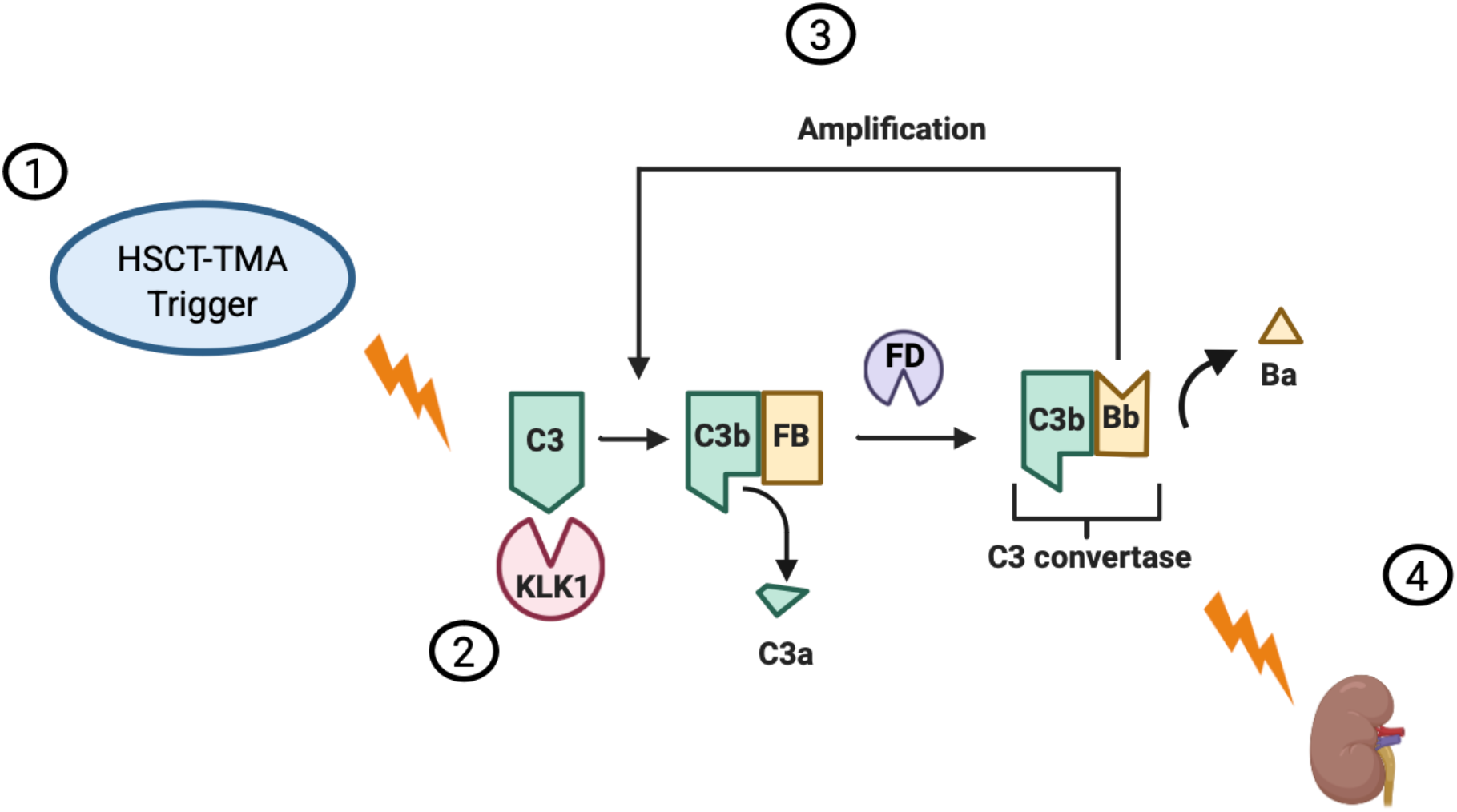

## Introduction

The complement system is a humoral component of the innate immune response, comprising plasma proteins that react nonspecifically to foreign entities. Activation occurs through three proteolytic cascades – the classical, lectin, and alternative pathways – which converge on the cleavage of complement factor C3. This generates anaphylatoxins, opsonins, and membrane attack complexes that mediate pathogen clearance and immune modulation, while inactivation mechanisms tightly regulate this response to mitigate collateral tissue destruction.^1^ Inflammatory stimuli shift this balance, enhancing complement activity that complicates disease outcomes and interventions.

The alternative pathway (AP) progresses to full activation when C3 is cleaved, generating cleavage products C3a and C3b. C3b binds factor B (FB), forming C3bB. The serine protease factor D (FD) then cleaves FB in C3bB to release Ba and form the C3 convertase (C3bBb) that further generates C3a and C3b by cleaving free C3, amplifying AP activation.^2–6^ When uninhibited, abundant C3b nonspecifically deposits onto cell surfaces and binds C3 convertases, forming the C5 convertases that activate the terminal complement pathway.

Dysregulation of the alternative complement pathway (AP), leading to uninhibited tissue destruction, has been recognized in many disease states,^7–11^ including the inflammatory thrombotic microangiopathies (TMAs). Hematopoietic stem cell transplant-associated TMA (HSCT-TMA) is a complement-mediated TMA that occurs after HSCT, usually in the setting of another inflammatory or immune process.^12–14^ The pathophysiologic mechanisms of HSCT-TMA are poorly defined but related to a complex interplay between inflammation, complement activation, and endothelial injury that leads to microvascular thrombosis and ultimately end organ damage, particularly of the kidneys.^15^ We and others have shown that the AP is overly activated in HSCT-TMA, as measured by generation of the FB cleavage product Ba.^16–18^ Various triggers occurring in the peri-transplant period have been associated with the development of HSCT-TMA, including inflammatory mediators and the calcineurin inhibitor class of immunosuppressants (such as cyclosporine A [CsA] and tacrolimus),^15,19–22^ but whether these triggers cause AP activation, or are simply associated with it, is unclear.

In contrast to the classical and lectin pathways that are activated upon stimulation, the AP is continuously activated at a low level through a tick-over mechanism. In this mechanism, a baseline level of hydrolyzed C3 (C3_H2O_) is continuously maintained in blood, priming a robust generation of C3b.^23,24^ However, formation of C3b by non-complement, inflammatory proteases^24–26^ suggests that mechanisms other than C3_H2O_ tick-over may contribute to AP activation. Tissue kallikreins – a family of 15 paralogous serine proteases that are structurally homologous to the serine protease domain of plasma kallikrein^27,28^ – are notably elevated in diseases that present with inflammation.^29,30^ In mice challenged with lipopolysaccharide (LPS), translation of tissue kallikrein 1 (*Klk1*) but not other detectable paralogs (*Klk5, Klk6*, and *Klk14*) is upregulated in the renal endothelium.^31^ Canonically, KLK1 cleaves kininogens to produce kinins, potent vasodilators with inflammatory effects,^27,28^ and is inhibited by the serine protease inhibitor (SERPIN), kallistatin.^32^ KLK1 exhibits trypsin- and chymotrypsin-like specificity^33^ resulting in broad substrate preferences.^34–37^ Because C3 is remarkably upregulated in inflamed glomerular microvascular endothelial cells (GMVECs) of the kidney,^38,39^ we hypothesized that KLK1 activates the AP by cleaving C3 to generate C3b, contributing to renal endothelial injury of HSCT-TMA. In this study, we demonstrate that C3 is a KLK1 substrate that yields a functional cleavage product, establishing an association between KLK1 dysregulation and AP overactivation in models of HSCT-TMA.

## Methods

### Patient Samples

Citrated patient plasma samples used in this study were obtained from a pediatric prospective longitudinal cohort study of all-comers to HSCT.^18^ HSCT-TMA study definition and sample collection, processing, and storage are as previously described.^18^ A subset of patients were randomly selected for our study. Samples collected pre-HSCT and at 60 and 90 days post-HSCT were analyzed. Patients without HSCT-TMA served as controls. This study was approved by the Institutional Review Board at Baylor College of Medicine.

### Assays for Human Ba and KLK1

#### Human Ba antigen

Ba was measured from citrated patient plasma by commercial ELISA kit according to the manufacturer’s instructions (Quidel, A033), as previously reported.^17^

#### Human KLK1 inactivation

To measure KLK1 activity in patient plasma, the amount of KLK1 in the KLK1/kallistatin complex was analyzed by semi-quantitative western blotting. Citrated patient plasma (1 µL) was resolved on a 4-15% SDS-PAGE (BioRad, 4561086) under reducing conditions. Proteins were transferred to Immobilon-FL PVDF membrane (Millipore, IPFL00010) and immunostained with rabbit anti-human KLK1 (1 µg/mL; Abcam, ab28289) followed by IRDye800CW donkey anti-rabbit IgG secondary antibody (50 ng/mL; LI-COR, 926-32213). The blots were imaged on a LI-COR Odyssey DLx and analyzed with LI-COR Image Studio (version 6.0) software. To assess KLK1 inactivation in pre-HSCT and post-HSCT samples, free-KLK1 (30 kD) and kallistatin-bound KLK1 in the KLK1/kallistatin complex (80 kD) was quantified and a ratio of bound-KLK1 to total (free + bound) KLK1 was calculated.

### Murine Studies

#### Mice

Adult male and female C57BL/6J mice (The Jackson Laboratory, 000664), 9-15 weeks old, were used for experimentation. Mice were bred and housed at Rice University BioScience Research Collaborative. All murine procedures and experiments were approved by the Rice University Institutional Animal Care and Use Committee.

#### Blood Drawing

To obtain plasma samples, whole blood was collected from anesthetized mice (isoflurane or ketamine/xylazine) from either the retro-orbital sinus with a 3.2% (w/v) buffered sodium citrate-filled capillary tube, or from the vena cava of mice pre-treated with buffered sodium citrate prior to blood collection with a 27G needle on a 1mL syringe. The ratio of sodium citrate to blood was 1:9 (v/v). Plasma was separated from the whole blood by centrifugation at 5000xg for 15 minutes.

#### Murine exposure to CsA + LPS

Mice were bled retro-orbitally at least one week prior to CsA + LPS injection to obtain a baseline sample (pre-CsA + LPS). WT mice were intraperitoneally injected with CsA (30 mg/kg; Tocris, 1101) and LPS (5 mg/kg; from *Escherichia coli* O111:B4, Sigma-Aldrich, L2630) or DMSO (Sigma-Aldrich, D1435) as control. Twenty-four hours after CsA + LPS injection, the mice were anesthetized with ketamine/xylazine, blood was collected (retro-orbital sample followed by terminal bleed through IVC), and kidneys were dissected.

#### Kidney preparation and staining for complement deposition and thrombomodulin (TM) shedding

Dissected kidneys were immersed in 10% buffer formalin for 24 hours and subsequently subjected to a sucrose gradient (10-30% sucrose in phosphate buffered saline [PBS], w/v) prior to optimal cutting temperature (OCT) compound embedding, cryosectioning (2 µm slices), and affixing to microscope slides by the Baylor College of Medicine Center for Comparative Medicine Comparative Pathology Laboratory. The frozen kidney sections were permeabilized with 0.2% Triton X-100 in PBS, treated with TrueBlack Lipofuscin Autofluorescence Quencher (Biotum, 23007), and blocked with 5% normal donkey serum (Jackson ImmunoResearch Laboratories, 017-000-121). These kidney sections then were stained with either biotinylated goat anti-mouse C3d antibody (20 µg/mL; R&D Systems, BAF2655) and streptavidin Alexa Fluor 488 conjugate (5 µg/mL; Invitrogen, S32354), following streptavidin/biotin blocking according to manufacturer’s instructions (Vector Laboratories, SP-2002) to block endogenous biotin and streptavidin binding sites, or goat anti-mouse TM antibody (10 µg/mL; R&D Systems, AF3894) and donkey anti-goat IgG Alexa Flour 488 (5 µg/mL; Invitrogen, A-11055). Cell nuclei were stained by 4’,6-diamidino-2-phenylindole (DAPI) included in the mounting medium (Vector Laboratories, H-1800). After mounting, the stained kidney sections were examined by immunofluorescence microscopy using a Nikon Diaphot TE300 microscope equipped with a 60x objective, SensiCamQE CCD camera (Cooke Corp.), motorized stage and dual filter wheels (Prior) with single band excitation and emission filters for FITC/TRITC/CY5/DAPI (Chroma). Images were acquired and processed using IP Lab software version 3.9.4r4 with a fluorescence colocalization module (Scanalytics, Inc.).

### Assays for Murine KLK1, Ba, C3d, soluble TM (sTM), and Creatinine

#### Semi-quantitative immunoblotting for KLK1 inactivation, Ba, and C3d

KLK1 was assessed by measuring the amount of KLK1 bound in the KLK1/kallistatin complex in mouse plasma samples. Mouse plasma (0.5 µL) was resolved and blotted as above. For KLK1 inactivation, blots were stained with sheep anti-mouse KLK1 (R&D Systems, AF7928)/ donkey anti-sheep IgG (H+L) Alexa Fluor 680 (Invitrogen, A-21102). For murine Ba, blots were immunostained with a rabbit anti-mouse FB antibody (1 µg/mL; Pacific Immunology, custom design) that recognizes FB and Ba, followed by IRDye 800CW goat anti-rabbit IgG secondary antibody (50 ng/mL; LI-COR, 926-32211). For murine C3d, blots were immunostained with a biotinylated goat anti-mouse C3d antibody (0.1 µg/mL; R&D Systems, BAF2655) that recognizes C3 and C3d, followed by IRDye 800CW Streptavidin (0.1 µg/mL; LI-COR, 926-32230). Immunostained blots were imaged above. Murine KLK1 inactivation was quantified as for human KLK1 inactivation (**Supplemental Figure 1A**). Murine Ba formation was quantified as a ratio of signal intensities between cut FB (Ba) and total FB (FB + Ba, **Supplemental Figure 1B**). Murine C3d levels were determined as a ratio of signal intensities between C3d and total C3 (C3 + C3d, **Supplemental Figure 1C**).

#### Murine sTM Levels

sTM was measured from citrated plasma by commercial ELISA kit according to the manufacturer’s instructions (R & D Systems, MTHBD0).

#### Murine Creatinine Levels

Creatinine levels were measured from deproteinized citrated plasma using an enzymatic assay according to the manufacturer’s instructions (Crystal Chem, 80350). Plasma was deproteinized using 10K MWCO centrifugal devices (Cytiva, OD010C34) according to manufacturer’s instructions.

### Fluid phase studies

#### Cleavage of AP components by KLK1

Recombinant human KLK1 (200 µg/mL; R&D Systems, 2337-SE-010) was incubated for 1 hour with thermolysin (2 µg/mL; Promega, V4001) in activation buffer (25 mM Tris, pH 7.4, 50 mM NaCl, 5 mM CaCl_2_) at 37°C for proteolytic activation (rhKLK1a). C3 (0.29 µM; Complement Technologies, A113) alone or mixed with FB (0.11 µM; Complement Technologies, A135) was incubated with buffer (25mM Tris, pH 7.4, 150 mM NaCl, 10mM MgCl_2_, ± 5 mM CaCl_2_, ± 0.005% Tween-20), thermolysin (65 ng/mL), non-activated rhKLK1 (0.20 µM), or rhKLK1a (0.20 µM) at 37°C. Samples of the reactions were taken at indicated times and resolved on 4-15% SDS-PAGE (BioRad, 4561086) under reducing conditions. The gels were blotted onto Immobilon-FL PVDF membrane (Millipore, IPFL00010) and immunostained with goat anti-C3 (1:200,000 dilution; Complement Technologies, A213)/IRDye680RD donkey anti-goat IgG secondary antibody (50 ng/mL; LI-COR,926-68074) ± rabbit anti-FB (Pacific Immunology, custom design)/IRDye800CW donkey anti-rabbit IgG secondary antibody (50 ng/mL; LI-COR, 926-32213) to assess for C3 and FB cleavage products, respectively. The blots were imaged on a LI-COR Odyssey DLx and analyzed with LI-COR Image Studio (version 6.0) software.

#### C3b function: Ba generation by KLK1-generated C3b

To generate C3b, purified C3 (0.34µM) was incubated with rhKLK1a (1.2µM) for 90 min at 37°C for complete cleavage of C3 to C3b (rhKLK1a-generated C3b). As negative controls, purified C3 was incubated with equivalent amounts of thermolysin or activation buffer found in the rhKLK1a-generated C3b reaction. The rhKLK1a-generated C3b (0.29 µM), or purified C3b (0.29 µM; Complement Technologies, A114), was incubated with FB (0.11 µM) and FD (0.003 µM) at 37°C for 10 min and subsequently stored at 4°C for 24 hours until Ba was measured by ELISA (Quidel, A033). Reactions without C3b (i.e., FB + FD and rhKLK1a + FB + FD) were included as negative controls.

### Data Analysis

Data were analyzed using GraphPad Prism 10. Mean and standard deviation (SD) were calculated and significance determined by unpaired or paired t-tests, or one-way ANOVA with Tukey’s multiple comparisons test as indicated. P-values <0.05 were considered significant. Pearson correlation coefficients were calculated and statistically tested to determine strength of associations.

## Results

### HSCT-TMA patients demonstrated increased AP activation and decreased KLK1 inactivation

We analyzed AP activation and KLK1 inactivation from a subset of plasma samples obtained from a prospectively enrolled pediatric HSCT cohort at Texas Children’s Hospital, wherein enrolled patients were longitudinally screened for HSCT-TMA with serial plasma collection pre- and post-HSCT.^18^ From this cohort, 11 patients who developed HSCT-TMA were included in our analysis; 7 were female, the median age was 12 years (range 3-21), the median time to diagnosis was 79 days after HSCT, and all received calcineurin inhibitors for immunosuppression. 19 patients who did not develop HSCT-TMA (controls) were included in our analysis of AP activation, and 10 of these were included in our analysis for KLK1 inactivation. Of the 19 control patients, 6 were female, the median age was 8 years (range 2-20) and all were immunosuppressed with a calcineurin inhibitor.

We confirmed our earlier observation that the AP is activated in HSCT-TMA, using Ba as a marker.^17,18^ In control patients who underwent HSCT but did not develop TMA, Ba levels remained stable between pre- and 60-90 days post-HSCT samples (668 ± 240 ng/mL vs. 745 ± 264 ng/mL, respectively; *n*=19, *p*>0.3; **Fig. 1B**). In contrast, patients who developed HSCT-TMA showed significantly elevated Ba levels in the peri-diagnosis period compared to pre-HSCT (1,688 ± 909 ng/mL vs. 762 ± 317 ng/mL, respectively; *n*=11, *p*<0.01; **Fig. 1A**).

**Fig. 1.**
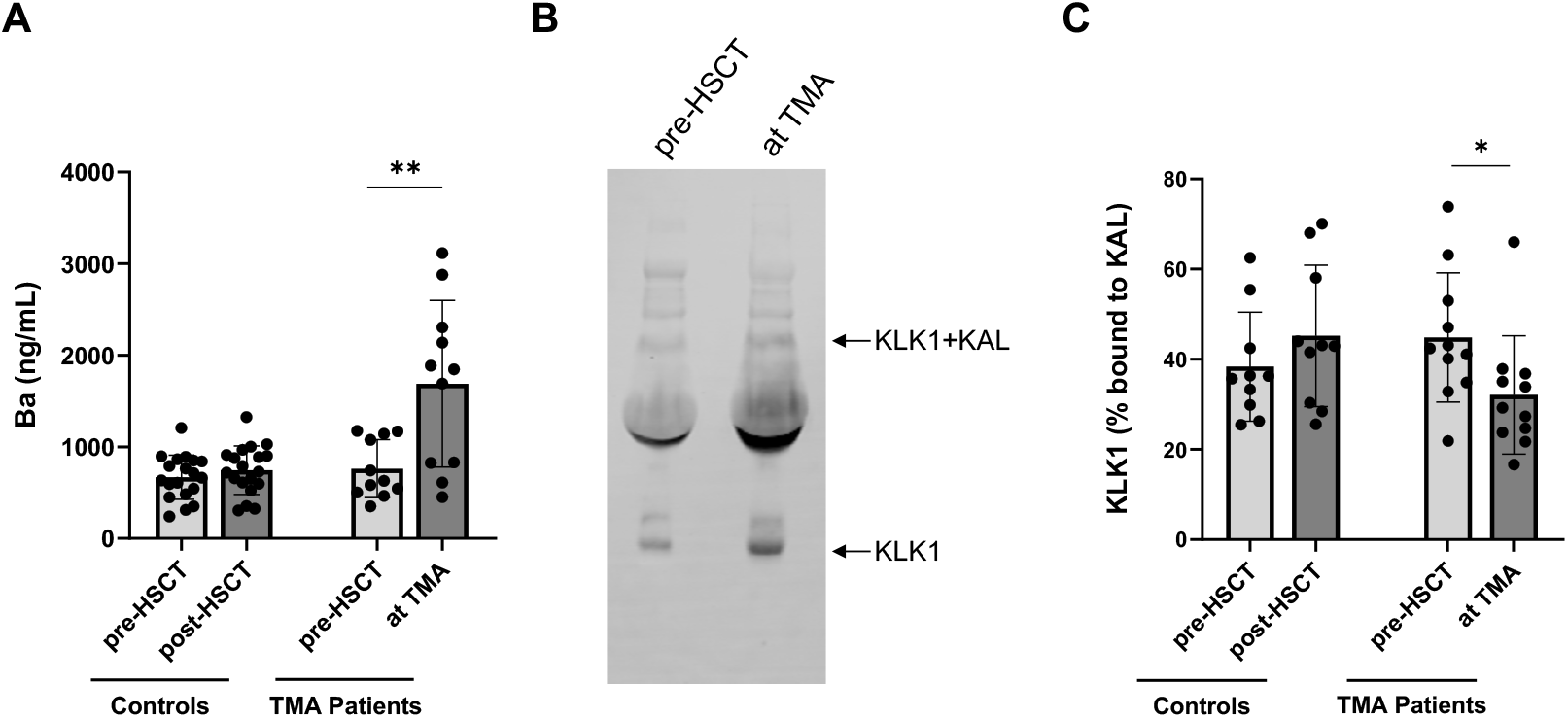
AP activation and KLK1 inactivation in patients with HSCT-TMA. Ba levels and KLK1 inactivation were measured in pediatric HSCT patient samples obtained prior to and after HSCT. **(A)** Ba levels (by ELISA) in control patients (HSCT patients without TMA) before and at a time period of 60-90 days after HSCT (n=19), and in HSCT-TMA patients before HSCT and at diagnosis of TMA (median 79 days, n=11). **(B and C)** KLK1 inactivation was measured by fluorescent immunoblotting. **(B)** Representative blot of the free-KLK1 and KLK1 + KAL complex in a patient with HSCT-TMA. Free (~30 kD) and KAL-bound (~80 kD) KLK1 was quantified to calculate the bound fraction relative to the total (free + bound). **(C)** Amount of bound KLK1 in the KLK1 + KAL complex in control patients before and at a median of 90 days post-HSCT (n=10), and in patients with HSCT-TMA before HSCT and at diagnosis of TMA (n=11). Mean ± SD was calculated and significance was determined with a paired t-test. * = *p*<0.05, ** = *p*<0.01.

We subsequently evaluated plasma levels of inactivated KLK1 by semiquantitative immunoblotting for the KLK1/kallistatin complex (**Fig. 1B, representative blot**). In patients who did not develop HSCT-TMA, the relative amount of inactivated KLK1 was comparable between post-HSCT and pre-HSCT (45.2 ± 15.7% vs. 38.4 ± 12.1%, respectively; n=10, *p*>0.1, **Fig 1C**). In contrast, significantly less KLK1 was found in complex with kallistatin among HSCT patients diagnosed with TMA relative to pre-HSCT (32.1 ± 13.1% vs. 44.8 ± 14.4%, respectively; *n*=11, *p*<0.05; **Fig. 1C**), consistent with reduced KLK1 inactivation during disease onset.

### Diminished KLK1 inactivation coincided with AP activation, renal complement deposition, renal endothelial injury, and renal dysfunction in mice exposed to HSCT-TMA triggers

To investigate the impact of HSCT-TMA triggers on KLK1 activity and AP activation, we measured plasma levels of KLK1 inactivation, Ba, and C3d in C57BL/6J mice challenged with 30 mg/kg CsA and 5 mg/kg LPS. Consistent with KLK1 inactivation in patients who progressed to HSCT-TMA, less KLK1 was found to be in complex with kallistatin (i.e., reduced KLK1 inactivation) in mice after a single challenge with HSCT-TMA triggers compared to before (30.2 ± 19.8% vs. 56.1 ± 25.6%, respectively, n=6, *p*<0.01, **Fig. 2A**, immunoblots in **Supplemental Fig. 1A**). Similarly consistent, the combinantion of CsA and LPS led to increased AP activation as indicated by greater FB proteolysis (6.3 ± 6.1% [pre-challenge] vs. 20.6 ± 9.6% [post-challenge, n=6, *p*<0.05], **Fig. 2B, Supplemental Fig. 1B**). Furthermore, C3d (a C3 activation product downstream of C3b generation) was higher in mice after challenge with CsA and LPS compared to pre-challenge levels (22.5 ± 5.3% vs. 9.0 ± 0.6%, respectively, n=6, *p*<0.01, **Fig. 2C, Supplemental Fig. 1C**).

**Fig. 2.**
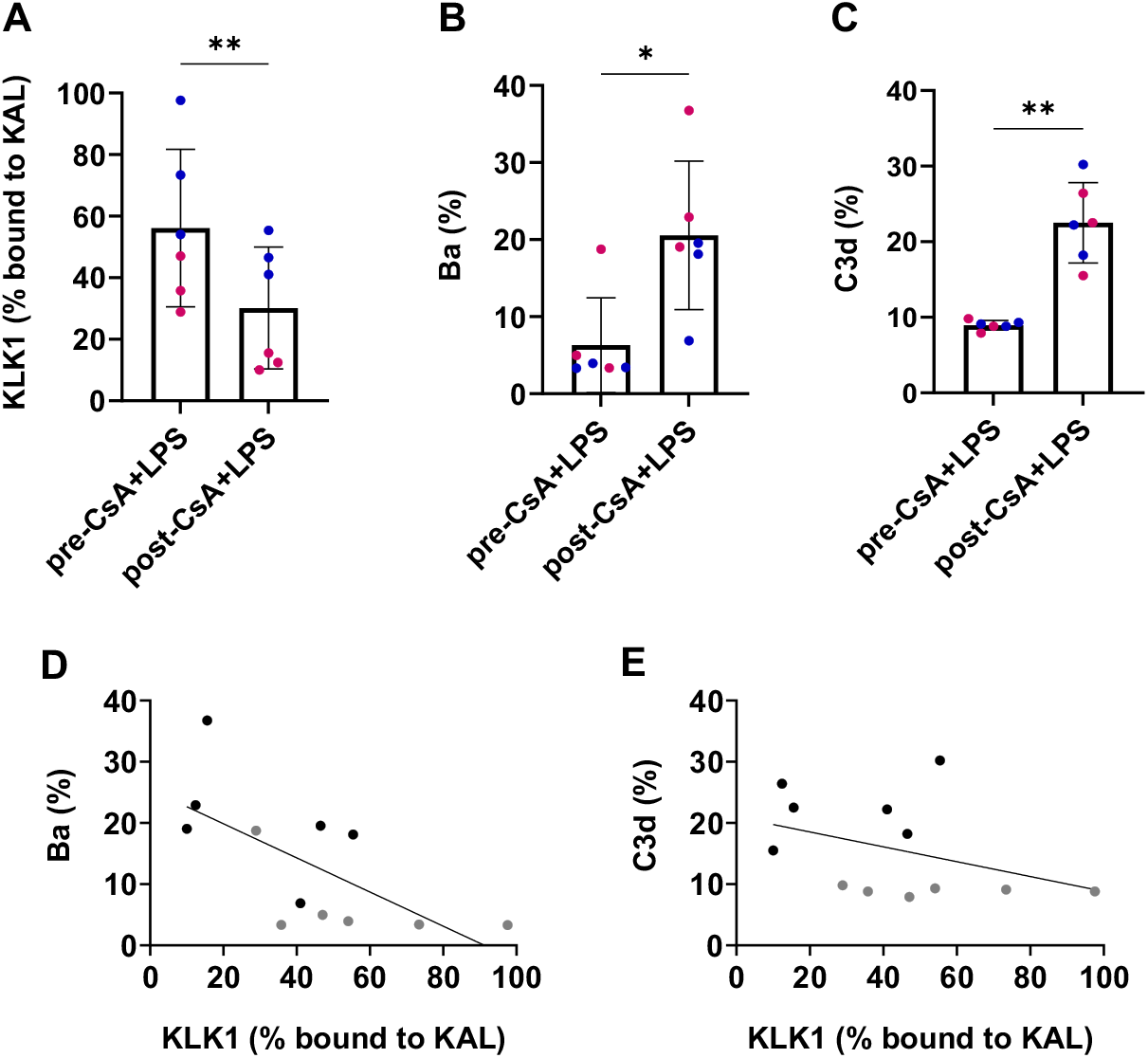
KLK1 inactivation and AP activation in mice exposed to CsA and LPS. Adult C57BL/6J mice were injected intraperitoneally with CsA (30 mg/kg) + LPS (5 mg/kg). Citrated plasma was collected pre- and 24 hours post-CsA+LPS injection and analyzed by western blotting for KLK1, Ba, and C3d (C3b cleavage product); see Supplemental Figure 1. **(A)** Free and KAL-bound KLK1 were quantified and the bound fraction relative to the total (free + bound) was calculated. **(B)** Ba and FB were quantified and the percentage of Ba to total FB (FB + Ba) was calculated. **(C)** C3d and C3 were quantified and percentage of C3d to total C3 (C3 + C3d) was calculated. Columns = Mean, Error bars = SD, Blue = male, Pink = female. Significance was determined with a paired t-test. * = *p*<0.05, ** = *p*<0.01. **(D)** Ba and **(E)** C3d negatively correlated with KLK1 (Pearson correlation coefficient, r = −0.6684, *p*<0.05; Pearson correlation coefficient, r = −0.3914, *p*>0.2; respectively). **(D and E)** Gray = pre-CsA + LPS-injected mice, Black = 24 hours post-CsA + LPS-injected mice.

To assess the relationship between KLK1 inactivation and AP activation, we performed Pearson correlations. We found a significant, negative correlation between KLK1 inactivation and Ba levels (r=-0.6684, *p*<0.05, **Fig. 2D**), suggesting an inverse association. While also an inverse relationship, a correlation between KLK1 inactivation and C3d level was not statistically detectable (r=-0.3919, *p*>0.2, **Fig. 2E**). Although these correlations seem inconsistent, the lack of a significant association between KLK1 inactivation and plasma C3d levels may also reflect a loss of circulating C3d to tissue deposition.

To assess the effects of complement activation on renal endothelial injury in the setting of HSCT-TMA triggers, kidneys were examined by immunohistochemistry for AP activation (deposition of C3d) and endothelial injury (TM shedding)^40–42^ 24 hours after challenge with 30 mg/kg CsA and 5 mg/kg LPS. C3d was deposited in higher amounts on the glomeruli of the challenged mice (**Fig. 3C**) compared to control (DMSO) mice (**Fig. 3A**), consistent with AP activation. Conversely, TM on the glomeruli surface was lost in mice challenged with CsA + LPS (**Fig. 3D**) compared to control mice (**Fig. 3B**), consitent with endothelial injury. Further supporting endothelial injury, sTM (TM that is shed into the plasma) was elevated in mice challenged with CsA and LPS compared to those challenged with DMSO (50.4 ± 14.5 ng/ml vs. 18.6 ± 0.9 ng/ml, respectively; n=3, *p*<0.05, **Fig. 3E**). To assess renal function in the setting of endothelial injury, creatinine was measured in both groups and was found to be higher in mice challenged with the HSCT-TMA triggers compared to those challenged with vehicle alone (0.36 ± 0.08 mg/dL vs. 0.06 ± 0.06 mg/dL, respectively; n=3, *p*<0.01, **Fig. 3F**), consistent with renal dysfunction. Pearson correlation showed a significant, positive correlation between sTM and creatinine levels (r=0.8331, *p*<0.05, **Fig. 3G**), suggesting an association between endothelial injury and renal dysfunction. Taken together, these data suggest that HSCT-TMA triggers may stimulate increased KLK1 activity and AP activation, resulting in renal complement deposition, renal endothelial injury, and impaired kidney function.

**Fig. 3.**
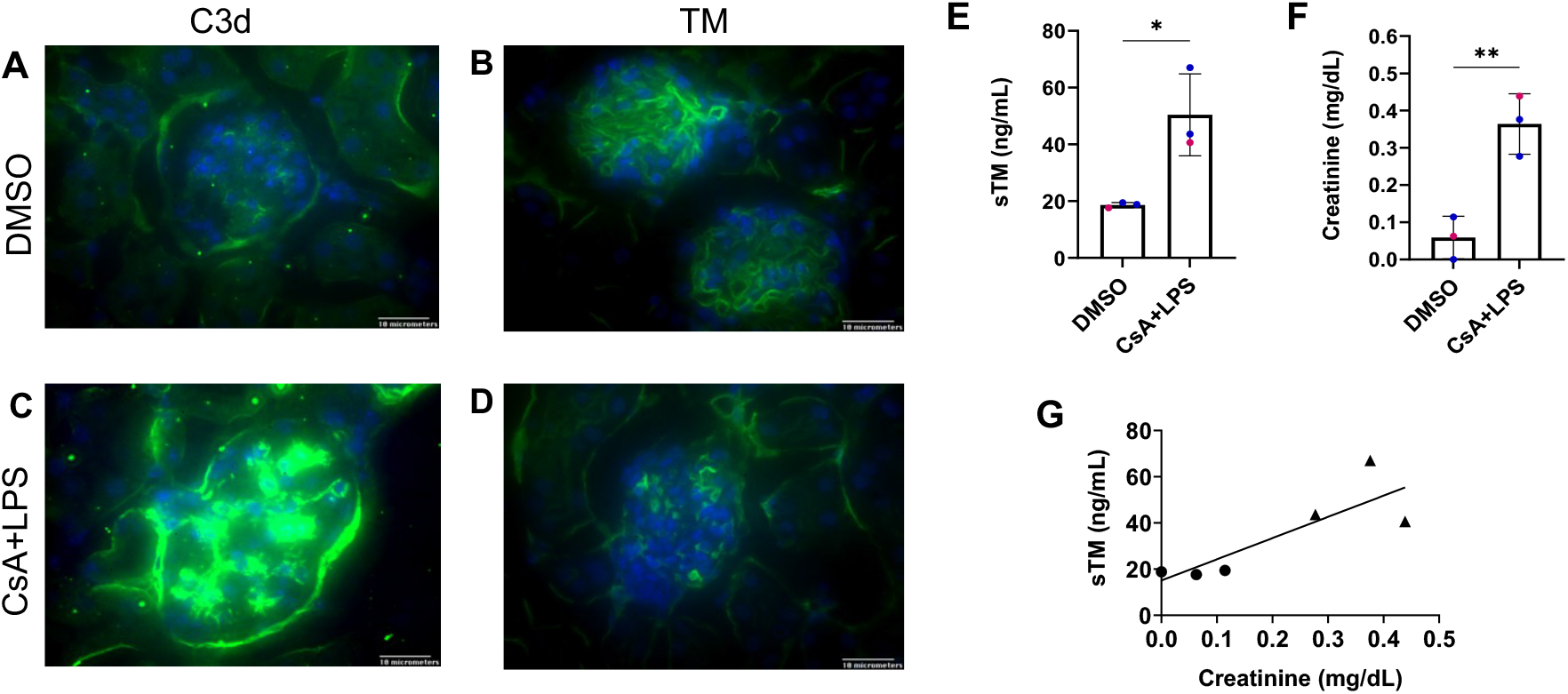
Renal complement deposition, endothelial injury, and dysfunction in mice exposed to CsA and LPS. Adult C57BL/6J mice were injected intraperitoneally with CsA (30 mg/kg) + LPS (5 mg/kg) or DMSO (control). Citrated plasma and kidneys were collected 24 hours post-injection. Mounted fixed frozen kidney sections were incubated with biotinylated anti-mouse C3d (C3b cleavage product)/streptavidin Alexa Fluor 488 (green, **A** and **C**) or anti-mouse TM/anti-goat IgG Alexa Flour 488 (green, **B** and **D**). Cell nuclei were stained with DAPI (blue). Stained kidney sections were imaged by immunofluorescence microscopy with a 60x objective. Scale bar = 10 micrometers. Plasma sTM was measured by ELISA **(E)**, and circulating creatinine was measured by enzymatic assay **(F)**. Columns = Mean, Error bars = SD, Blue = male, Pink = female. Significance was determined with an unpaired t-test. * = *p*<0.05, ** =*p*<0.01. **(G)** sTM positively correlated with creatinine in DMSO- (●) and CsA + LPS-treated (▲) mice. Pearson correlation coefficient, r = 0.8326, *p*<0.05.

### KLK1 cleaved C3 in the fluid phase

We tested for a direct relationship between KLK1 and C3 cleavage by incubating purified components together and assessing for AP cleavage products generated over time (up to 120 minutes). Activated, recombinant KLK1 (rhKLK1a) cleaved C3 by 20 minutes but did not cleave FB at any time point as noted by the absence of the 30 kD Ba fragment by immunoblotting (**Fig. 4A**). The C3-FB mixture incubated alone or with the KLK1 activator thermolysin remained uncut, demonstrating specific cleavage of C3 by rhKLK1a. To further characterize the rate of C3 cleavage by KLK1, rhKLK1a or non-activated KLK1 (i.e., incubation of KLK1 without thermolysin) was incubated together with C3, and the C3b generated over 5 to 120 minutes was quantified by immunoblotting (**Fig. 4B**). Whereas C3 cleavage was absent with non-activated KLK1, C3 was specifically cleaved by rhKLK1a within 5 minutes and increasingly over time, resulting in an observed second order rate constant 7.8×10^2^ ± 2.8×10^2^ M^−1^s^−1^ (**Fig. 4C**).

**Fig. 4.**
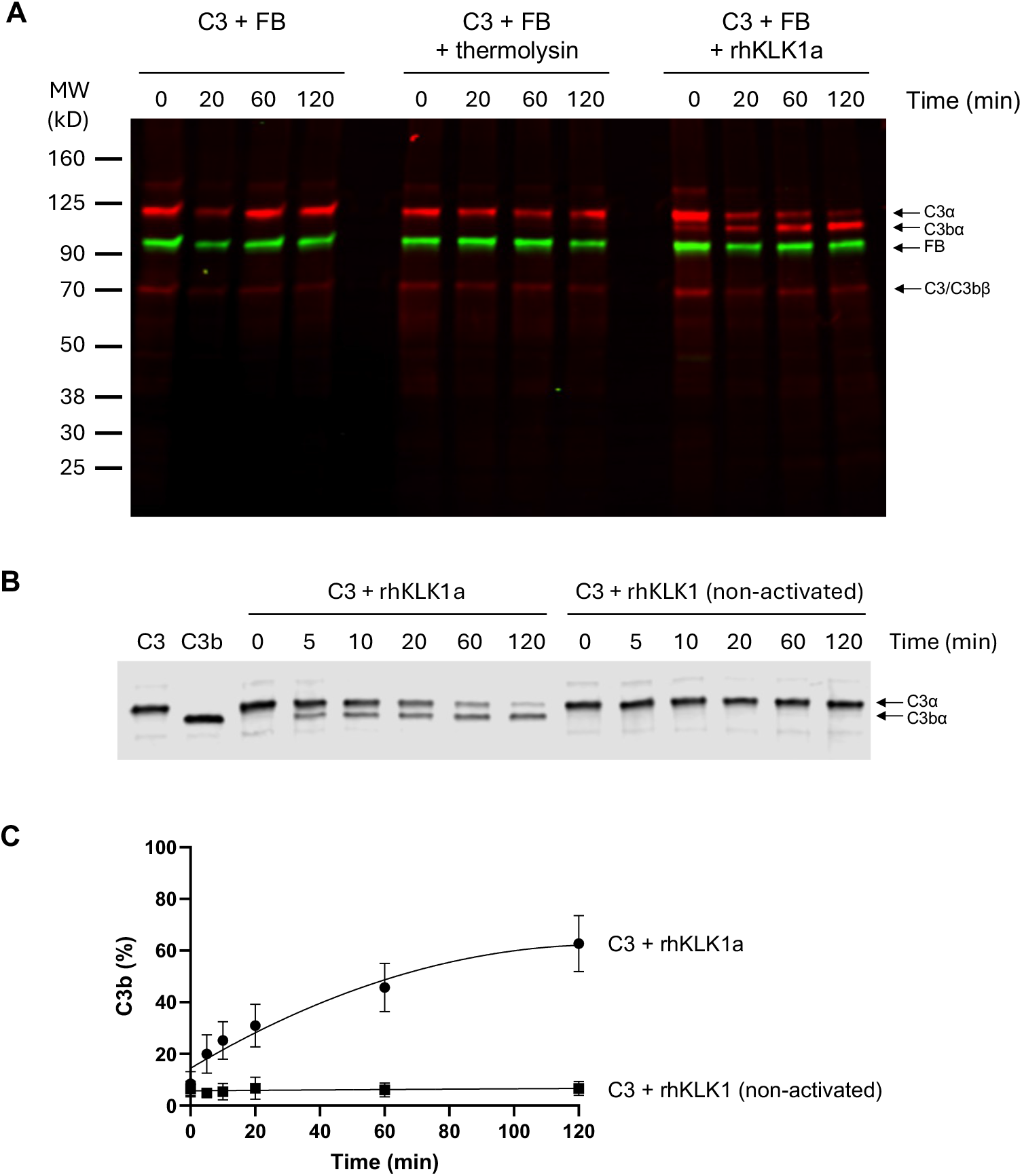
KLK1 cleavage of AP components. rhKLK1 was proteolytically activated by incubation with thermolysin (rhKLK1a). **(A)** C3 and FB were incubated with or without rhKLK1a (or thermolysin as a control) in buffer without Tween-20. Samples were taken at 0, 20, 60, and 120 min, resolved by SDS-PAGE under reducing conditions, and immunoblotted for the presence of C3 and FB cleavage products (C3b and Ba, respectively). C3 and its cleavage products were detected with goat anti-C3/IRDye680RD donkey anti-goat IgG secondary antibody (red); FB and its cleavage product, Ba, were detected with rabbit anti-FB/IRDye800CW donkey anti-rabbit IgG secondary antibody (green). C3 is composed of two disulfide-linked chains – alpha and beta. Upon cleavage, C3a is released from the alpha chain resulting in the generation of C3b (C3b*α* and C3b*β*). Proteolysis of FB yields cleavage product Ba (~30 kD). **(B)** C3 was incubated with activated and non-activated rhKLK1 in buffer containing Tween-20. Samples were taken at 0, 5, 10, 20, 60, and 120 min and resolved by SDS-PAGE under reducing conditions and immunoblotted for C3 cleavage products as described above. Purified C3 and C3b alone (two leftmost lanes) served as reference. **(C)** The amount of C3b*α* generated was quantified and calculated as a percent of total C3 (C3+C3b), n = 3. Fitted lines are to aid visualization.

### KLK1-cleaved C3b activates the AP by generating FB cleavage product Ba

To determine the activity of KLK1-generated C3b, we measured the amount of Ba produced upon incubation with FB and FD. In contrast to thermolysin or the absence of a protease, C3 was completely cleaved by excess rhKLK1a as indicated by immunoblotting (**Fig. 5A**). Incubation of the fully cleaved KLK1-generated C3b with FB and FD for 10 minutes generated significant amounts of Ba in comparison to controls (1,372 ± 23 ng/mL vs. FB + FD = 117 ± 6 ng/mL, *p*<0.0001, or vs. FB + FD + rhKLK1a = 188 ± 14 ng/mL, *p*<0.0001, respectively; n=3 per group, **Fig. 5B**). Likewise, plasma-derived C3b incubated with FB and FD generated Ba levels comparable to that of KLK1-generated C3b (1,353 ± 79 ng/mL vs. 1,372 ± 23 ng/mL, respectively; *p*=0.94), indicating that C3b generated by KLK1 is functionally similar to native C3b as a cofactor for FB proteolysis by FD. Consistent with Fig. 4A, rhKLK1a did not cleave FB into Ba in the presence of FD, further supporting the specific cofactor role of C3b.

**Fig. 5.**
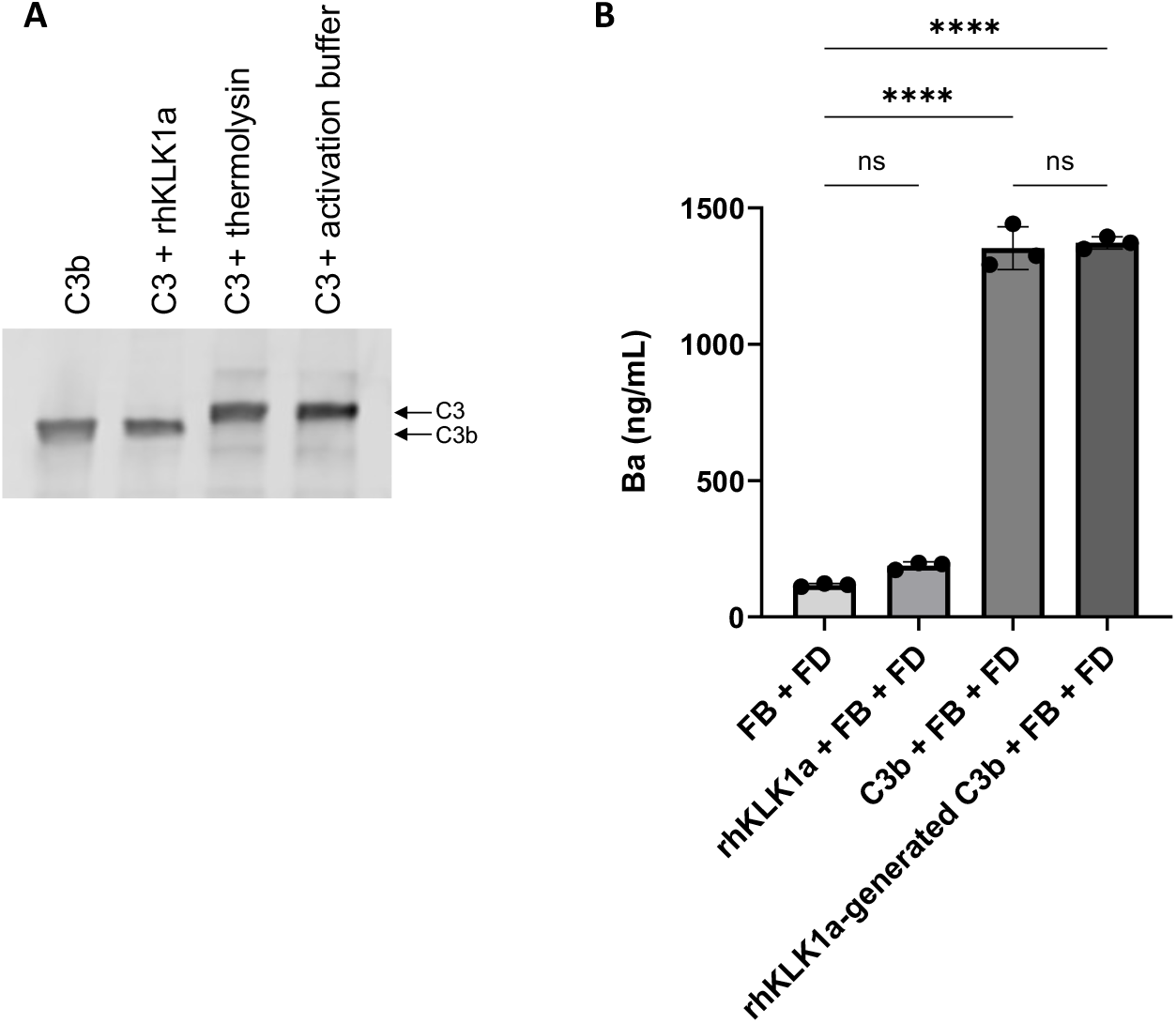
Ba generation by KLK1-generated C3b. **(A)** Purified C3 was incubated with rhKLK1a at 37°C for 90 minutes to ensure complete cleavage into C3b. Purified C3 was also incubated with thermolysin or activation buffer, which are both present in the rhKLK1a preparation. Reaction samples were resolved and immunoblotted for C3 cleavage products, which were compared to purified C3b to verify complete cleavage. n=3 for each reaction (representative blot shown). **(B)** Purified C3b or rhKLK1a-generated C3b was incubated with FB + FD at 37°C for 10 min, followed by storage at 4°C for 24 hours. FB + FD and rhKLK1a + FB + FD served as controls. Ba generated in the reactions was measured by ELISA. n=3 for each reaction. Mean ± SD was calculated and significance determined by one-way ANOVA with Tukey’s multiple comparisons test. **** *p* <0.0001.

## Discussion

The outcomes of HSCT-TMA are historically poor, with mortality reaching as high as 50%, along with significant renal morbidity.^19,43,44^ These poor outcomes are secondary to diagnostic challenges, lack of early recognition due to overlap with many other post-HSCT related conditions, and inadequate treatment modalities. The best currently available treatment for HSCT-TMA, the anti-C5 monoclonal antibody eculizumab, targets the terminal complement pathway (TP) and has shown limited efficacy: studies report that one third of patients treated with eculizumab have no response, and 25-50% have long term kidney impairment.^45,46^ These inadequate responses to anti-C5 therapy suggest that other mechanisms of complement dysregulation may underlie the progression of HSCT-TMA.

In this study we evaluated the potential for the serine protease KLK1 to cleave C3 as a mechanism for AP activation in HSCT-TMA. With a cohort independent from our previous report,^17^ we confirm that HSCT-TMA presents with increased circulating Ba levels, and we now demonstrate that inactivation of KLK1 by kallistatin is diminished in these patients. Similar findings with our mouse model suggest that immunosuppression and inflammation may have a significant role in complement-mediated HSCT complications that include renal endothelial injury and impaired kidney function.

Hypertension is a common complication of calcineurin inhibition and TMA more broadly. Unmasking of complement defects in TMA associated with severe and malignant hypertension (without HSCT, calcineurin inhibition, or hematologic symptoms) points to a link between complement overactivation and renal vasculopathy.^47,48^ Dysregulated interactions between the renin-angiotensin, kallikrein-kinin, and complement systems may give rise to a spectrum of vascular and thromboinflammatory syndromes that have overlapping features.^49^ Renin, an aspartic acid protease that cleaves angiotensin to induce vasoconstriction, has been demonstrated to cleave C3 into functional components of the AP.^50^ However, renin is expressed in 10-fold lower amounts than KLK1 in the inflamed kidney^31^ and was reported to have little-to-no pathophysiological impact in complement disorders.^51^

Others have reported C3 as a substrate for plasma kallikrein,^52–54^ a serine protease of the contact (intrinsic) pathway of coagulation that is secreted as a zymogen (prekallikrein), requiring activation by activated factor XII (FXIIa) to become functional.^55,56^ However, renal transplant patients immunosuppressed with CsA have comparable levels of FXIIa as healthy controls.^57^ In contrast to plasma kallikrein, KLK1 is intracellularly processed and secreted as an active enzyme.^28,49^ Rats treated with CsA have upregulated expression of intrarenal KLK1, associated with KLK1-mediated hypertension.^58^ The profundity of renal injury in HSCT-TMA, the robustness of KLK1 translation over other organs (brain, heart, liver, and lung),^31^ and C3 cleavage by KLK1 suggest that local responses to systemic inflammation and immunosuppression may underlie pathologies that develop into fulminant HSCT-TMA.,

Although debated,^24^ current understanding posits that C3 is hydrolyzed at a low, constant rate into C3(H_2_O), a C3b-like cofactor for FB proteolysis by FD to initiate AP activation. Rapid decay of the C3(H_2_O) convertase via dissociation or by complement inhibitors (e.g., factors H and I) is thought to protect against AP amplification.^24,59,60^ The slow observed kinetics of C3b generation by KLK1 in comparison to the catalytic efficiency of the C3 convertase, C3bBb (3.1 x 10^5^ M^−1^s^−1^),^61^ suggests an initiating role (instead of amplification) for KLK1 in AP activation. Functionally indistinguishable from native C3b, accumulation of KLK1-generated C3b may be sufficient to ignite AP amplification. With decreased KLK1 inactivation by kallistatin and inadequate AP inactivation in HSCT-TMA,^16,62^ we propose that inflammation and immunosuppression in HSCT shifts the balance between C3b generation and inactivation towards AP amplification as part of a complex disease process that involves KLK1 and results in endotheliopathy.

Understanding the pathophysiology of AP overactivation in HSCT-TMA, as well as in other poorly defined inflammatory TMAs, would pave the way for novel therapies that target the excessive cleavage of complement components without compromising immune surveillance. Our data demonstrate a role for KLK1 in AP activation via C3 cleavage and support an association between KLK1, AP activation, endothelial injury, and renal dysfunction in HSCT-TMA. Further study is needed to determine the function of the KLK1/kallistatin axis in preventing or dampening complications of HSCT or other inflammatory microvascular diseases.

## Acknowledgments

We thank the patients for their contributions. We thank Dr. Audrey Cleuren (Oklahoma Medical Research Foundation) for her assistance with reviewing differential translation of endothelial *Klk1* in mice.

Funding was provided by the National Institutes of Health (AI175828, SES, AY; HL154688, AY; HL159271, AL; 3OT2OD032581-01S1, AL), American Society of Hematology Scholar Award (SES, AY, and AL), Alex’s Lemonade Stand Foundation Innovation Grant (SES and AY), American Cancer Society Discovery Boost Grant (SES and AY), Mary R. Gibson Foundation (SES and AY), CPRIT Scholar in Cancer Research, supported by Cancer Prevention and Research Institute of Texas (RR190104, AL), and Career Development Award from Conquer Cancer, the ASCO Foundation (AL).

## Author Contributions

SES and AY conceived the project. SES, PMJ, and AY designed the experiments. SES, MR, JB, SB, and AL collected and provided patient samples. SES, PMJ, VQ, and BL conducted the experiments. SES, PMJ, and AY analyzed and interpreted the results. SES, PMJ, and AY wrote this report.

## Figure Legends

**Supplemental Fig. 1.**
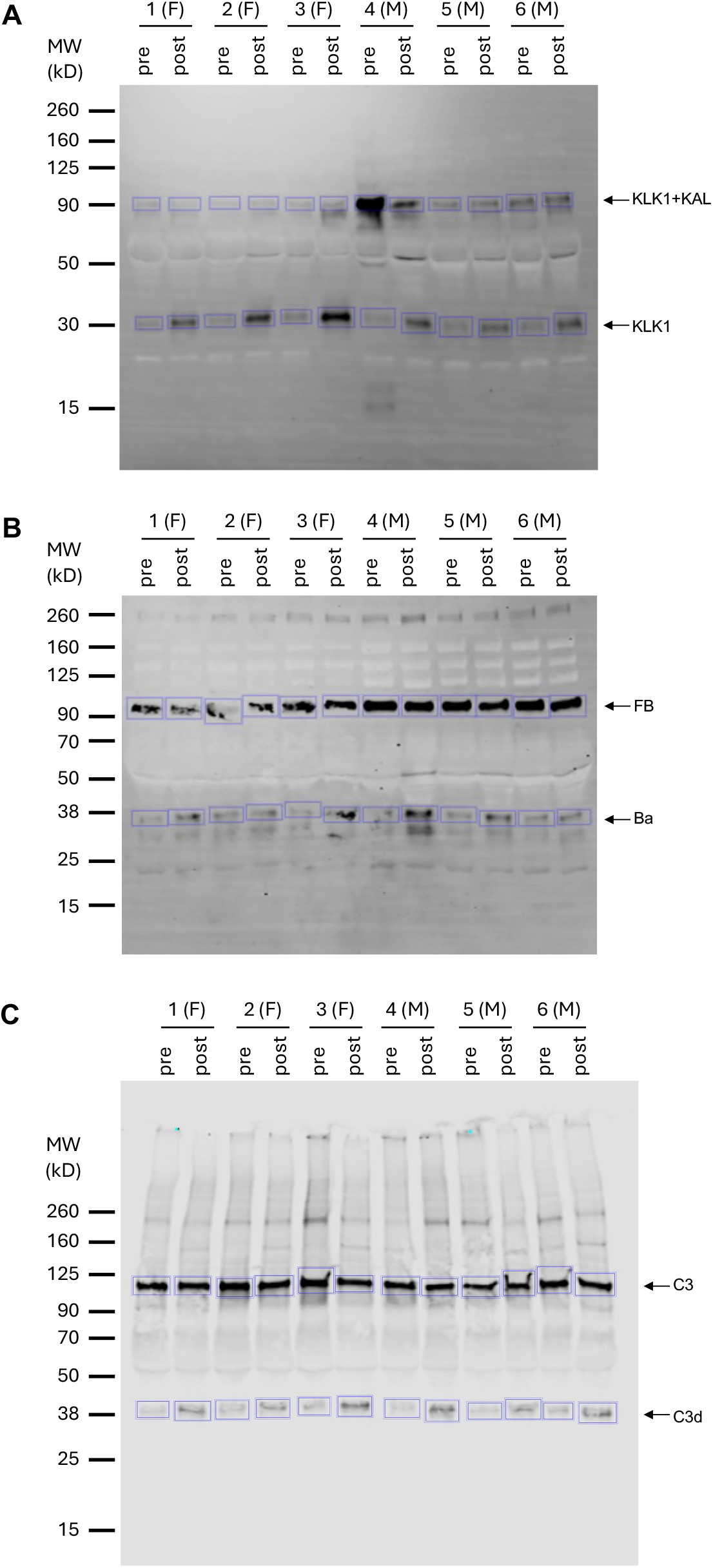
Western blots used for KLK1, Ba, and C3d calculations in Fig. 2. Citrated plasma (1 µL) from adult C57BL/6J mice injected with CsA + LPS (pre-injection [pre] and 24 hours post-injection [post]) was resolved on 4-15% SDS-PAGE under reducing conditions and immunoblotted for **(A)** KLK1 and KLK1+KAL, **(B)** FB and Ba, and **(C)** C3 and C3d. **(A)** Free and KAL-bound KLK1, **(B)** Ba and FB, and **(C)** C3d and C3 were quantified. Blue boxes indicate the quantified areas. MW = Molecular weight marker. F = female mouse, M = male mouse.

